# Targeting the TR4-induced RCC cells-derived exosomally initiated signaling increases Sunitinib efficacy

**DOI:** 10.1101/2022.12.07.519440

**Authors:** Zhenwei Wang, Yin Sun, Huiyang Xu, Chi-Ping Huang, Bo Cheng, Fuju Chou, Bosen You, Xiaofu Qiu, Guosheng Yang, Chawnshang Chang

**Affiliations:** Department of Urology, Guangdong Second Provincial General Hospital, Guangzhou, 510317, China; George Whipple Lab for Cancer Research, Departments of Urology and Pathology and The Wilmot Cancer Institute, University of Rochester Medical Center, Rochester, NY, 14642 USA; Sex Hormone Research Center and Department of Urology, China Medical University/Hospital, Taichung, 404, Taiwan; Department of Urology, the Affiliated Hospital of Southwest Medical University, Luzhou, 646000, China; Department of urology, the Second Affiliated Hospital of Harbin Medical University, Harbin 150086, China; Shanghai East Hospital, Tongji University School of Medicine, Shanghai, 310000, China

**Keywords:** Renal cell carcinoma, TR4, Angiogenesis, Exosomes, Sunitinib

## Abstract

Sunitinib is the first-line therapy for metastatic clear cell renal cell carcinoma (RCC) via suppressing neoangiogenesis and tumor growth. The detailed mechanisms, especially whether and how RCC cells can impact endothelial cells sensitivity to Sunitinib, remain unclear. Here, we found that TR4 was commonly upregulated in RCC tissue and relative to tumor angiogenesis. Tube formation and Mouse aortic ring assay showed that the overexpression or knockdown of TR4 in RCC cells enhanced or reduced the angiogenesis of endothelial cells and their resistance to Sunitinib in vitro. Mechanistically, We found that TR4 transcriptionally increase ADAM15 expression, as a consequence, exosomes carrying relatively large amounts of ADAM15 secreted from RCC cell resulted in activating the EGFR phosphorylation and reducing the efficacy of Sunitinib in endothelial cells. Targeting the TR4-induced renal cancers-derived exosomelly initiated signaling with a small molecular, CRM197, increases sunitinib efficacy in vitro and xenograft tumor models. Taken together, our findings indicate a novel function of TR4 in RCC blunted the efficacy of sunitinib via exosomal ADAM15-induced activation of EGFR signaling pathway in endothelial cells.

## Introduction

Renal cell carcinoma(RCC), a main subtype of kidney cancer, is one of the most lethal urological tumors with an annual mortality rate of approximately 90,000 according to WHO statistics[1]. Approximately one-third of patients with RCC have metastases at presentation[2]. Sunitinib has been used as the first-line therapy to effectively suppress metastasis RCC in the first 6-15 months. However, most treatment may fail due to development of resistance to Sunitinib, and mechanisms for this remain unclear[3]. Therefore, studying the detailed mechanisms responsible for the therapy resistance, remains needed.

Sunitinib has antiangiogenetic effect through inhibition of VEGF signals in endothelial cells and platelet-derived growth factor receptor beta (PDGFR-β) expressed on the supporting pericytes[4]. Although Sunitinib also has a direct antitumor effect through inhibiting receptor tyrosine kinase KIT[5]and Fms-related tyrosine kinase 3 (Flt3)[6] in the disease where these signals are biologically relevant, no somatic mutations in target receptor tyrosine kinase (RTK) genes have been found in human RCC[7]. It is generally agreed that Sunitinib acts primarily through an antiangiogenic mechanism in addition to its ability to directly target RCC tumors[8]. As tumor angiogenesis is closely linked with tumor progression, we were interested in examining how RCC cells can impact the endothelial cell Sunitinib sensitivity.

The TR4 (testicular orphan receptor 4), a 67-kDa protein, belongs to the nuclear receptor superfamily[9]. Members of this family act as ligand-activated transcription factors and play roles in many biological processes such as development, cellular differentiation, and homeostasis[10]. Early studies indicated that TR4 have higher expression in RCC patients with lower recurrence-free survival (HR=1.587) than other patients and promoted RCC metastasis and vasculogenic mimicry(VM) formation *via* modulating HGF/MET functionand miR490-3p/VIMENTIN signals[11, 12]. Importantly, our early studies also indicated that a reduced TR4 activity augments the antitumor effects of sunitinib in xenograft tumor models via regulating sunitinib sensitivity of RCC cell lines[13]. Sunitinib as a multi-target inhibitor of tyrosine kinases is believed to be effective primarily through suppressing tumor angiogenesis as well as directly suppressing tumor proliferation. However, whether TR4 in RCC might also play a role in regulating endothelial cell sensitivity to Sunitinib have not been elucidated.

Exosomes are small (30-100 nm) membrane-encapsulated vesicles secreted by many cell types, including cancer cells. They are one means of intercellular communication among cancer cells and endothelial cells to regulate activities of cytokines[14] and growth factors. Although studies have been increasing in exosomes/microparticles from cancer cells to many other cell types such as macrophages [15], endothelial cells[16]and stromal cells [17], it is still unclear how cancer cells communicate with endothelial cells *via* exosomes.

The human A Disintegrin and Metalloproteinase 15 (ADAM15) is a member of the ADAM family of catalytically active disintegrin membrane metalloproteinases that function as proteolytic processing of cytokines, growth factors and adhesion molecules[18]. It has been shown that ADAM15 induces progression in several cancers including gastric[19], lung[20], colon[21]and prostate[22, 23]. ADAM15 as a mediator of angiogenesis is expressed in endothelial cells and up-regulated on the angiogenic endothelial cell surface[24, 25]. In addition, ADAM15 promotes cell proliferation, migration and sprouting of endothelial cells and smooth muscle cells[26]. However, the function of ADAM15 in RCC angiogenesis is still unclear.

This present study revealed that TR4 were significantly elevated in RCC patients and relative to tumor angiogenesis. Additionally, we demonstrated that TR4 in RCC may play an important role in activating the EGFR signaling pathway and reducing the efficacy of sunitinib in endothelial cell for angiogenesis through enhancing RCC-derived exosomes transferring ADAM15 between RCC cells and endothelial cells. We delineated a novel pathway for TR4’s effects on the efficacy of Sunitinib in RCC as well as provided a mechanistic explanation, and more importantly, we provided potential therapeutic approaches to overcome this resistance.

## Results

### Overexpression of TR4 is correlated with the progression and angiogenesis of RCC

Previously, we have found that the expression of TR4 results in a lower survival in RCC patients[12]. Furthermore, it could promote RCC VM formation and enhance the RCC resistance to Sunitinib. Whether TR4 can also alter Sunitinib resistance *via* other mechanisms remains unclear. In this study we found that the expression of TR4 was detected in normal renal cells and six different RCC cell lines. Western blot analysis indicated that TR4 is highly expressed in RCC cell lines(**Fig. 1A**). IHC analyses were used to study the expression of TR4 in RCC tissue and normal kidney tissue. The IHC results indicated that TR4 was significantly upregulated in 74.5%(38/51) of RCC tissues examined compared to that in 26.1%(6/23) of the normal kidney tissues (**Fig. 1B**). Moreover, by IHC analysis of TR4 and CD31 in 51 RCC clinical samples, we found that high TR4 expression was correlated with high microvascular density, suggesting that overexpression of TR4 might be related to tumor angiogenesis (**Fig. 1C**).

**Figure 1.**
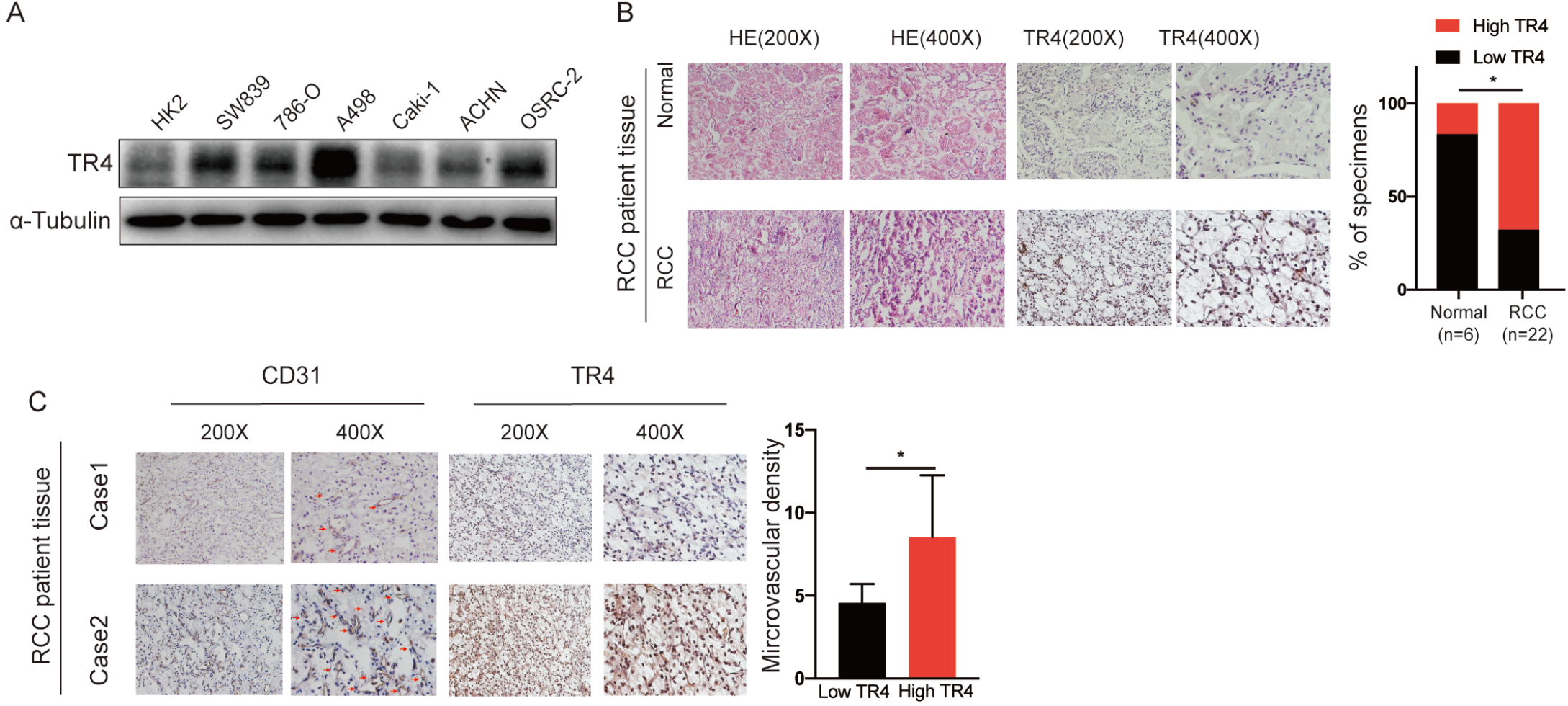
Overexpression of TR4 is correlated with the progression and angiogenesis of RCC. **A:**Expression of TR4 protein was detected in HK-2 and six different RCC cancer cell lines. **B:** IHC (HE) analysis of representative cases shows expression of TR4 protein in RCC tissues and adjacent normal kidney tissue (left) High vs low expression of TR4 in Normal (n=23) and RCC (n=51) specimens. **C:** IHC staining were used to detect the expression of TR4 and CD31 in human RCC tissues. Two representative cases are shown.

### TR4 expression in RCC cells alters angiogenesis and endothelial cell sensitivity to Sunitinib

As Sunitinib primarily suppresses tumor angiogenesis to suppress RCC progression[8], we were interested to see the potential impact of TR4 on the Sunitinib therapy.

We used the chamber co-culture system according to our previous reports[27]. We first co-cultured the RCC cell lines: including SW839, OSRC-2 and 786-O cells with human umbilical vein endothelial cells(HUVECs) and examined TR4 impact on the angiogenesis using tube formation assay with or without Sunitinib. The conditioned media (CM) from 72 hrs co-culture were collected and mixed with fresh media at 1:1 ratio to perform tube formation assasy with/without 1 μM Suntinib (see detailed outline in **Fig. 2A**).

**Figure 2.**
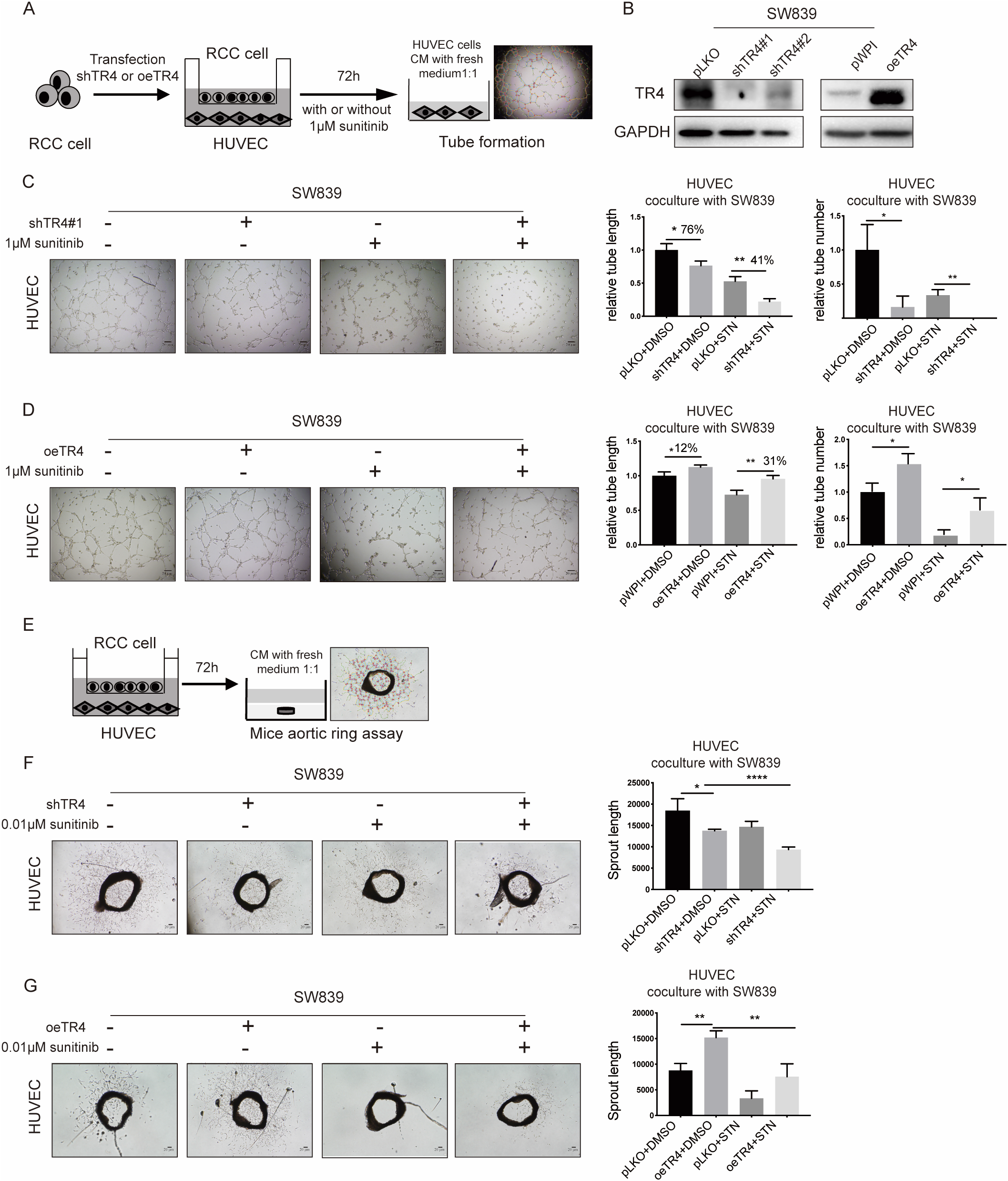
TR4 expression in RCC alters angiogenesis and endothelial cell sensitivity to Sunitinib. **A:** Outline of co-culture system. **B:** Western blot assay were perform to meassure TR4 in SW839 cells (+/-shTR4 and +/-TR4-cDNA). **C-D:**Tube formation assays were conducted in HUVECs cell co-cultured with SW839 cells transfected with (C) TR4-shRNA and pLKO with/without 1μM Suntinib (upper four images), or with (D) SW839 cells transfected with TR4-cDNA and pWPI with/without 1 μM Suntinib (lower four images). Tubular structures were photographed (150x) and quantified with image J software. **E:** Outline of Mouse aortic ring assay system. **F-G:** Mouse aortic ring assays were performed in the CM from SW839cells transfected with (F) TR4-shRNA and pLKOco-cultured system with/without 0.01 μM Sunitinib or (G) transfected with TR4-cDNA and pWPI co-cultured system with/without 0.01 μM Sunitinib. vessel sprouting from rat aorta was monitored by light microscopy and were Quantified with image J. For C, D, F, and G, quantitations are at the right of images.

The results revealed that CM from SW839 cells with knocked down TR4 via adding TR4-shRNA (**Fig. 2B**) have an decreasing the tube formation in HUVECs and increasing HUVEC sensitivity to Sunitinib treatment(**Fig. 2C**). Similar results were also obtained when we replaced SW839 cells with 786-O(Fig. S1A, C). In contrast, CM from SW839 cells with overexpressedTR4 *via* adding TR4-cDNA (**Fig. 2B**) have an increased capacity to promote tube formation in HUVEC cells (**Fig. 2D**). As expected, increasingTR4 in RCC SW839 cells also led to a decrease of HUVEC sensitivity to Sunitinib treatment (**Fig. 2D**). Consistent finding was observed in OSRC-2 cells (Fig. S1B, D).

We then applied a different approach with primary endothelial cells of mouse aorta [28]. The CM were used to assay the primary endothelial cells of mouse aorta with or without 0.01μM Sunitinib(**Fig. 2E, diagram**). The results confirmed the above findings, showing that CM from SW839 cells with shTR4 suppress the microvessel sprouting from the aortic ring and increased microvessel sensitivity to Sunitinib treatment(**Fig. 2F**), while CM from TR4 overexprssing SW839 cells could resulted in more quantity and length of vessels that sprouted from the aortic ring and decreased microvessel sensitivity to Sunitinib treatment (**Fig. 2G**).

These results suggested that TR4 in the RCC cells could promote the HUVEC angiogenesis and HUVEC cell sensitivity to Sunitinib treatment.

### TR4 in RCC cells impact the endothelial cell sensitivity to Sunitinib for angiogenesis via altering the exosomes to communicate between RCC cells and endothelial cells

We next explored the mechanism of how TR4 in RCC cells can impact the HUVEC cell sensitivity to the Sunitinib treatment, we first focused on classical secretory angiogenic factors including ANG1, CXCL8, DLL4, PGF, TGFbeta, VEGFA, EGF, and FGF-2 from RCC cells. However, the result show that overexpressing or knock down TR4 could not enhance or decrease the mRNA level of those genes (**Fig. S2A**)..

We then focus on the exosomes from RCC cells, as recent studies indicated that exosomes might play key roles in the cell-cell communication *via* exchanging certain bioactive factors, including membrane receptors, intracellular proteins, mRNAs, miRNAs, and/or organelles[29]. We first treated cells with the GW4869, an inhibitor of exosome biogenesis/release, to study its potential impact on the communication between RCC SW839 cells and endothelial HUVEC cells. The results revealed that treating with GW4869 led to partially reverse the RCC TR4-decreased endothelial HUVEC sensitivity to Sunitinib treatment(**Fig.3A**). We then isolated and examined the exosomes from the media of RCC cells using TEM assay[30]and found the size of exosomes from RCC SW839cells around100-200nm(**Fig. 3B**), consistent with known sizes of exosomes, as well as presence of CD9 and CD63, which are well known exosomes markers(**Fig. 3C**). Furthermore, results from tube formation assay revealed that adding exosomes from SW839-oeTR4 cells to HUVECs led to increase endothelial cells resistance to Sunitinib(**Fig. 3D**).

**Figure 3.**
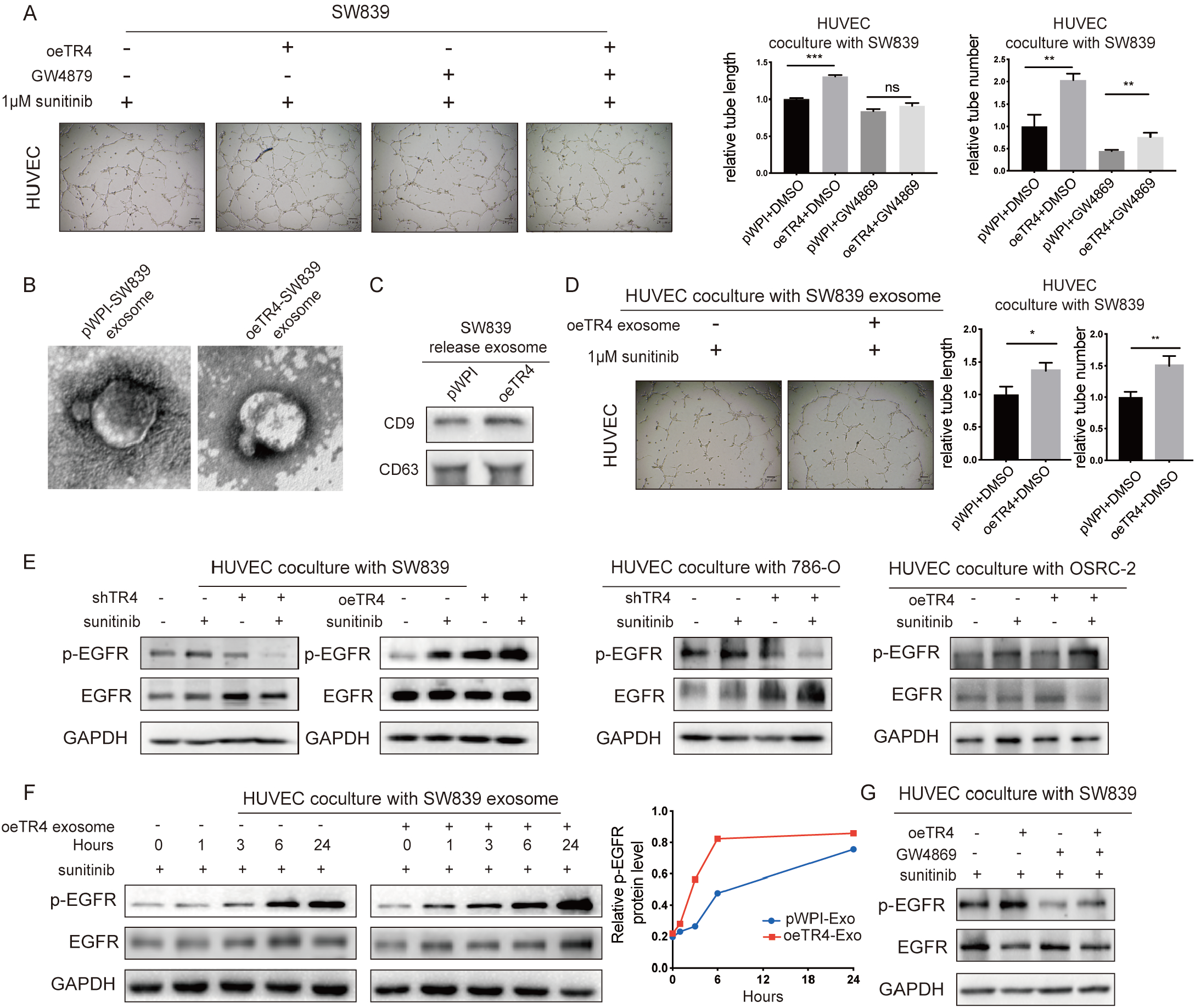
Mechanism dissection of how TR4 in RCC can impact endothelial cell sensitivity to Sunitinib: *via* releasing exosomes to communicate between RCC cells and endothelial cells. **A**. Tube formation assays were conducted in HUVECs co-culturedwith SW839 cells(+/-TR4-cDNA with 1 μM Sunitinib, +/-GW4869). **B**. Micrographs of exosomes isolated from SW839 cells transfected with pWPI (left) orTR4-cDNA (right). **C**. Western blot assays were performed to measure CD9 and CD63 in exosomes of SW839 cells (+/-TR4-cDNA). **D**. Tube formation assays in HUVEC cells with pWPI-SW839 exosomes or oeTR4-SW839 exosomes. **E**. Western blot assays were performed to measure phosphorylation and total EGFR in HUVECs co-cultured with SW839cells (+/-shTR4 and +/-1 μM Sunitinib or +/-TR4-cDNA and +/-1 μM Sunitinib) (left middle), or co-culturedwith 786-O cells(+/-shTR4 and +/-1 μM Sunitinib) (right middle), or co-culturedwith OSRC-2 cells (+/-TR4-cDNA and +/-1 μM Sunitinib) to test levels of phosphorylation and total EGFR. **F**. Western blot assays were performed to test protein levels of phosphorylation and total EGFR. Relative p-EGFR level at right. **G**. Western blot assay was performed to test protein levels of phosphorylation and total EGFR in HUVECs co-cultured with SW839 cells (+/-TR4-cDNA with 1 μM Sunitinib, +/-GW4869). Tubular structures were photographed (150x) and quantified with image J software.

Together, results from **Fig. 3A-D** suggest altering the TR4 in the RCC cells may influence the endothelial cell sensitivity to Sunitinib *via* altering the exosomes to communicate between RCC cells and endothelial cells.

### RCC exosomes can mediate the TR4 impact on endothelial cell sensitivity to Sunitinib treatment via inducing EGFR phosphorylation in endothelial cells

Next, to study how exosomes can mediate TR4’s impact on endothelial cell sensitivity to Sunitinib treatment, we focused on classical signaling pathways such as EGFR signaling on the endothelial cells. The results revealed adding CM from shTR4-SW839 and shTR4-786-O cells to HUVECs led to suppress EGFR phosphorylation which plays an important role during angiogenesis[31, 32] (**Fig. 3E**). Similarly, adding CM from oeTR4-SW839 and oeTR4-OSRC-2 cells to HUVEC led to activate EGFR phosphorylation(**Fig. 3E**). Consistent with this, exosomes from the oeTR4-SW839 RCC cells augmented EGFR phosphorylation in a more potent fashion(**Fig. 3F**), while GW4869 treatment could then prevent EGFR phosphorylation(**Fig.3G**).

These results suggest that TR4 in RCC cells may function *via* altering the exosomes to impact the EGFR activation, a critical signaling in endothelial cells angiogenesis, with a direct impact on the endothelial cell sensitivity to Sunitinib treatment.

### TR4 alter the exosomes content to impact endothelial cell sensitivity to Sunitinib treatment via increasing ADAM15 expression in RCC cells and exosomes

Next, we applied the GO class database[33] and PIPS database[34]analyses to identify 9 potential candidates whose function have been linked to the angiogenesis, exosomes functions, and upstream genes related to altering the EGFR signaling(**Fig. 4A**). Results from qPCR assay further found that shTR4in SW839 cells decreased the expression of ADAM15 (**Fig. 4B**). In contrast, oeTR4in SW839 cells increased the expression of ADAM15 (**Fig. 4B**).Moreover, results from western blot assay confirmed the regulation of ADAM15 expression in response to TR4 in SW839 cells (**Fig. 4C**).Significantly, Adam15 expression in the kidney of TR4 knockout mice[35] shows a much lower expression than that in the wildtype mice (**Fig. 4D**), indicating that TR4 regulates ADAM15 expression in a tissue-specific manner in the kidney.

**Figure 4.**
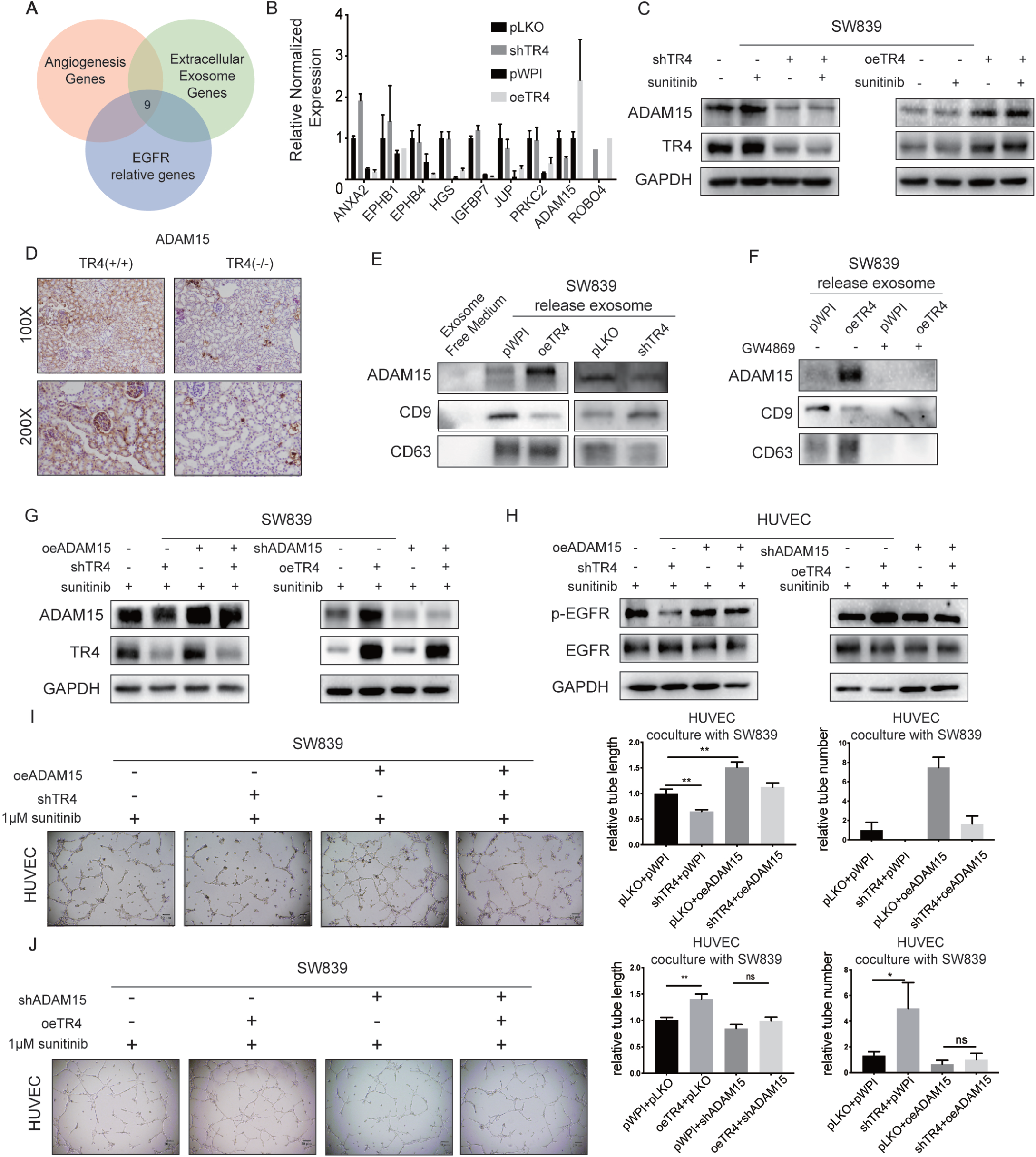
TR4 promoted angiogenesis and decreased Sunitinib sensitivity *via* altering ADAM15 expression. **A**. Diagram showing 9 potential candidate genes whose function areassociated with angiogenesis, exosomes functions, and upstream genes related to modulation of EGFR signaling. **B**. Real-time PCR assay for screening a set of 9 candidate genes in SW839 cells (+/-shTR4 and +/-TR4-cDNA). **C**. Western blot assay for ADAM15 expression in SW839 cells (+/-shTR4 and +/-1 μM Sunitinib or +/-TR4-cDNA and +/-1 μM Sunitinib). **D**. IHC staining was used to detect the expression of ADAM15 in normal mice and TR4 KO mice. **E-F**. Western blot assays for ADAM15 expression in exosomes from (E) exosomes-free FBS media (as control) and from SW839 cells (+/-shTR4 and +/-TR4-cDNA) (left panels), and in (F) exosomes from SW839 cells(+/-TR4-cDNA and +/-GW4869). **G**. Western blot assay for ADAM15 expression in SW839 cells (+/-shTR4 and +/-ADAM15-cDNA) with 1 μM Sunitinib (left panels) and SW839 cells (+/-TR4-cDNA and +/-shADAM15) with 1 μM Sunitinib (right panels). **H**. Western blot assays for phosphorylation and total EGFR expression in HUVECs co-culture with SW839 cells (+/-shTR4 and +/-ADAM15-cDNA) with 1 μM Sunitinib (left panels) and with SW839 cells (+/-TR4-cDNA and +/-shADAM15) with 1 μM Sunitinib (right panels). **I-J**. Tube formation assays were performed in HUVECs co-cultured with SW839cells (+/-shTR4 and +/-ADAM15-cDNA) with 1 μM Sunitinib (upper images) and (+/-TR4-cDNA and +/-shADAM15) with 1 μM Sunitinib (lower images). Tubular structures were photographed (150x) and quantified with image J software. Relative tube lengths are in middle panels and Relative tube numbers are in right panels.

Consistent with that, we also found oeTR4 in RCC cells led to release the exosomes that contain more ADAM15, suggesting increased TR4 increased the ADAM15 in RCC cells as well as in exosomes and decreased TR4 resulted in a decrease of ADAM15 (**Fig. 4E**), while GW4869 could block release of exosomes (**Fig. 4F**).

As expected and as shown in **Fig. 4G-H**, TR4-shRNA-decreased phosphorylation of EGFR in HUVECs was also partially reversed/blocked *via* increasing the ADAM15 expression in SW839 cell. In contrast, oeTR4-increased-phosphorylation of EGFR in HUVECs was partially reversed/blocked *via* decreasing the ADAM15 expression in SW839 cells. Furthermore, the interruption approach revealed that the TR4-shRNA-increased HUVECs sensitivity to Sunitinib treatment, were partially reversed/blocked by increasing the ADAM15 expression in SW839 cells (**Fig. 4I**). In contrast, oeTR4-increased tube formation and oeTR4-decreasedHUVECs sensitivity to Sunitinib treatment were partially reversed/blocked by suppressing the ADAM15 expression in SW839 cells(**Fig. 4J**). Interestingly, results from tube formation assay revealed that adding exosomes from SW839 cells to HUVECs have the similar result with condition medium (**Fig. S2B, C**).

Together, results from **Fig.4A-J** and **Fig. S2B-C**, suggest that TR4 in RCC cells may function *via* increasingADAM15 expression in RCC and exosomes to alter the angiogenesis/tube formation in endothelial cells and their sensitivity to Sunitinib treatment.

### TR4 transcriptionally increase ADAM15 expression

Since TR4 can increase ADAM15 expression at both protein and mRNA levels **(Fig. 4B-C)**, we were interested to see if TR4 can regulate ADAM15 mRNA expression *via* transcriptional regulation. Results from the chromatin immunoprecipitation (ChIP) assay with anti-TR4 antibody revealed that TR4 could weakly bind to the TR4 response-element (TR4RE)-III on the promoter region of ADAM15 in SW839 cells(**Fig. 5A-C**). therefore, we constructed an ADAM15 promoter luciferase reporter by inserting a 1 kb 5’ promoter region ofADAM15 containing TR4RE-III into the pGL3 luciferase backbone as well as a version with the mutated TR4RE-III**(Fig. 5D)**. Results from luciferase assay with the plasmid containing the ADAM15 gene 1 kb 5’promoter region with TR4RE show that TR4 can increase luciferase activity of wildtype, and not mutated form of the construct in SW839 cells (**Fig. 5E-F**). The above results indicated that TR4 may transcriptionally regulate the ADAM15 expression.

**Figure 5.**
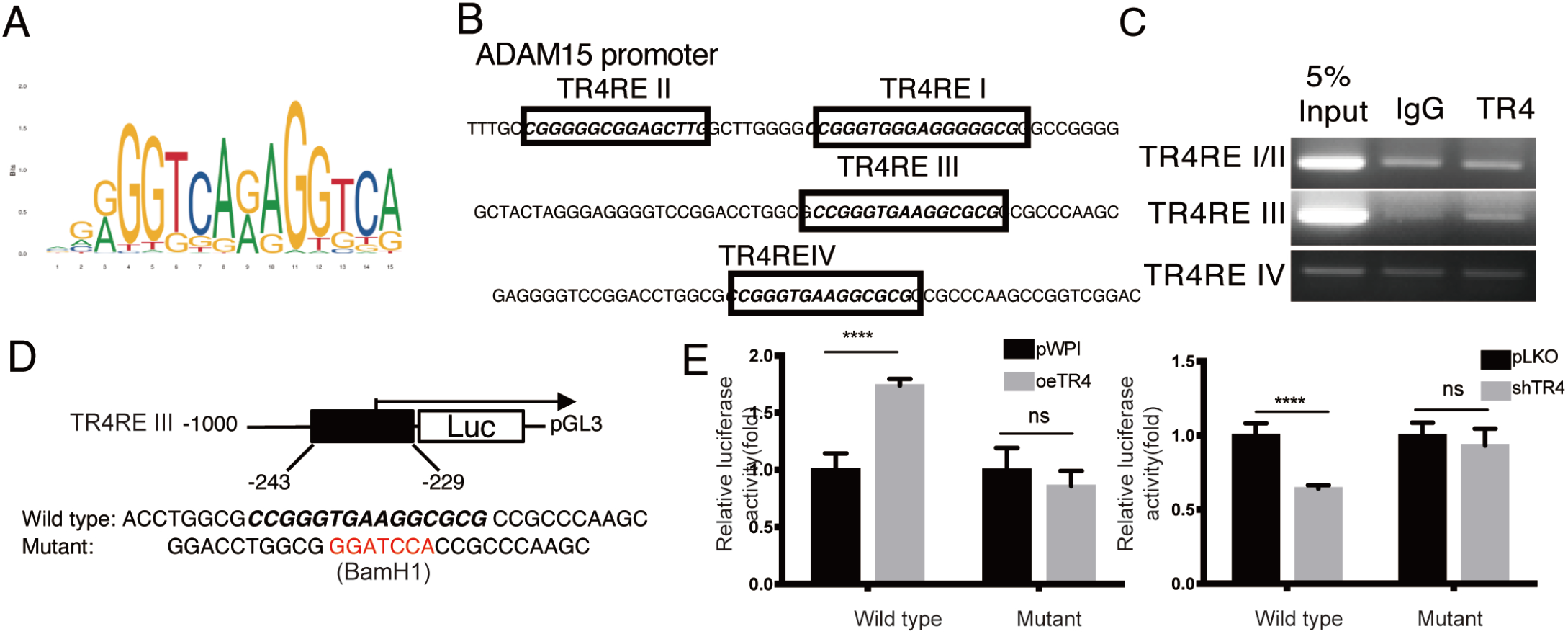
TR4 transcriptionally increase ADAM15 expression. **A**. TR4RE motif sequences. **B**. Four putative TR4RE within the 1-Kb promoter region of ADAM15 was predicted by JASPAR. **C**. ChIP assay DNAs were amplified by PCR reaction. **D**. Diagram of cloning the 1-kb ADAM15 promoter into pGL3 basic luciferase report vector (pGL3). **E**,**F**. Cotransfection of TR4RE wild type or mutant ADAM15 promoter pGL3-Luciferase constructs into SW839 with/without shTR4 or oeTR4. The luciferase assay was applied to detect the promoter activity. Data are presented as Mean ± SD.****, P < 0.0001; ns=not significant compared with the controls.

**Figure 6.**
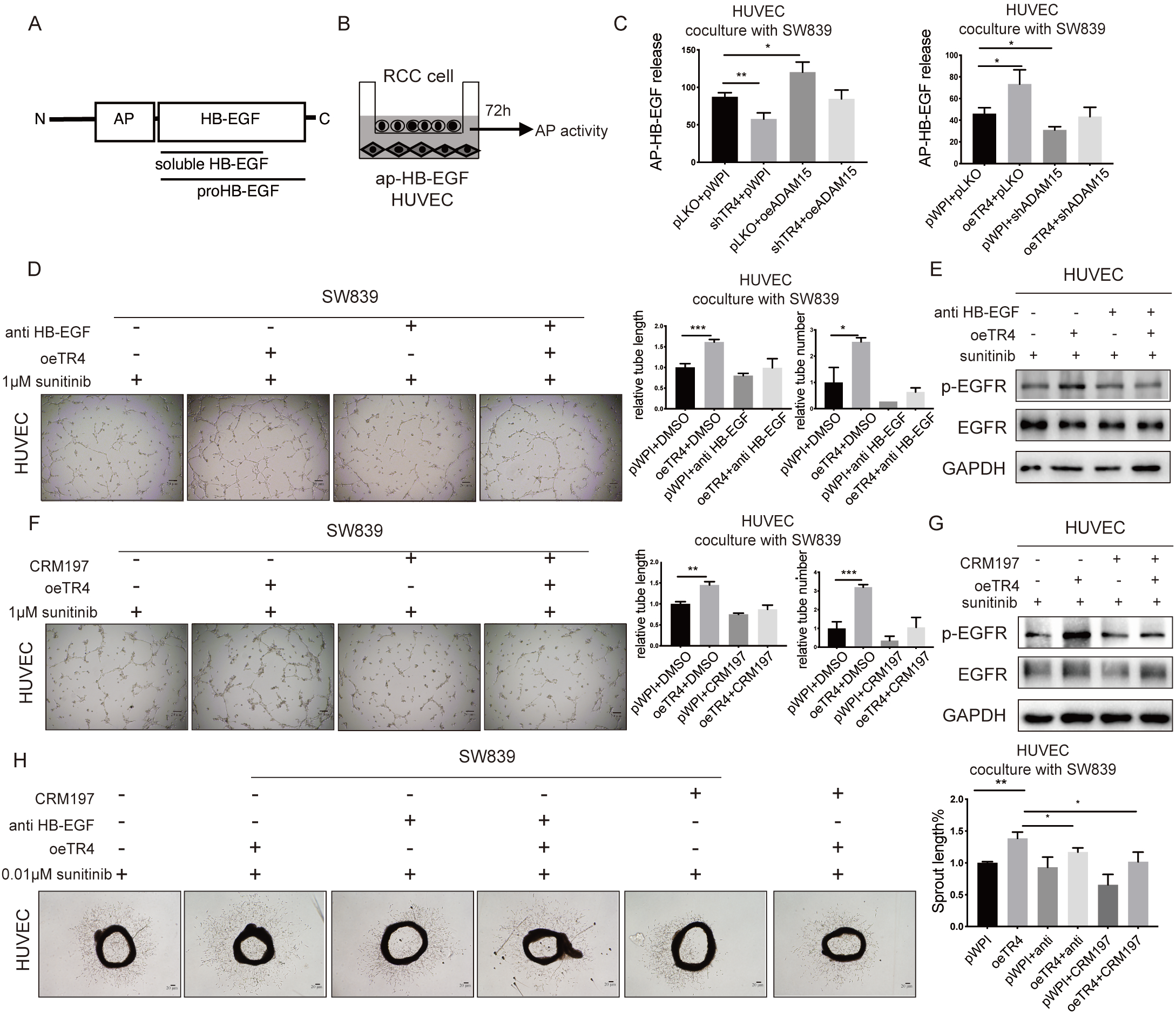
Mechanism dissection of how TR4/ADAM15 axis can alter angiogenesis in HUVEC cells: *via* increasing HB-EGF/EGFR signaling. **A**. Schematic representation of the AP–HB-EGF construct. **B**. Outline of alkaline phosphatase activity(AP-activity) system. **C**. AP-activity in CM from HUVEC cells co-cultured with SW839 cells (+/-shTR4 and +/-ADAM15-cDNA) (left panels) and (+/-TR4-cDNA and +/-shADAM15) (right panels). **D**. Tube formation assays in HUVEC cells co-cultured with SW839 transfected with TR4-cDNA and pWPI with/without HB-EGF neutralizing antibody. **E**. Western blot assays was performed to test protein levels of phosphorylation and total EGFR in HUVEC cells co-cultured with SW839 cells transfected with TR4-cDNA and pWPIwith/without neutralizing antibody. **F**. Tube formation assays in HUVEC cells co-cultured with SW839 cells transfected with TR4-cDNA and pWPI with/without CRM197. **G**. Western blot assay were performed to test protein levels of phosphorylation and total EGFR in HUVEC cells co-culture with SW839 cells(+/-TR4-cDNA and +/-CRM197) with 1 μM Sunitinib. **H**. Mouse aortic ring assay were performed in the CM from SW839cells transfected with TR4-cDNA and pWPI co-cultured system with 0.01 μM Sunitinib with/without neutralizing antibody or CRM197. Vessel sprouting from rat aorta was monitored by light microscopy and were Quantified with image J. Tubular structures were photographed (150x) and quantified with image J software.

**Figure 7:**
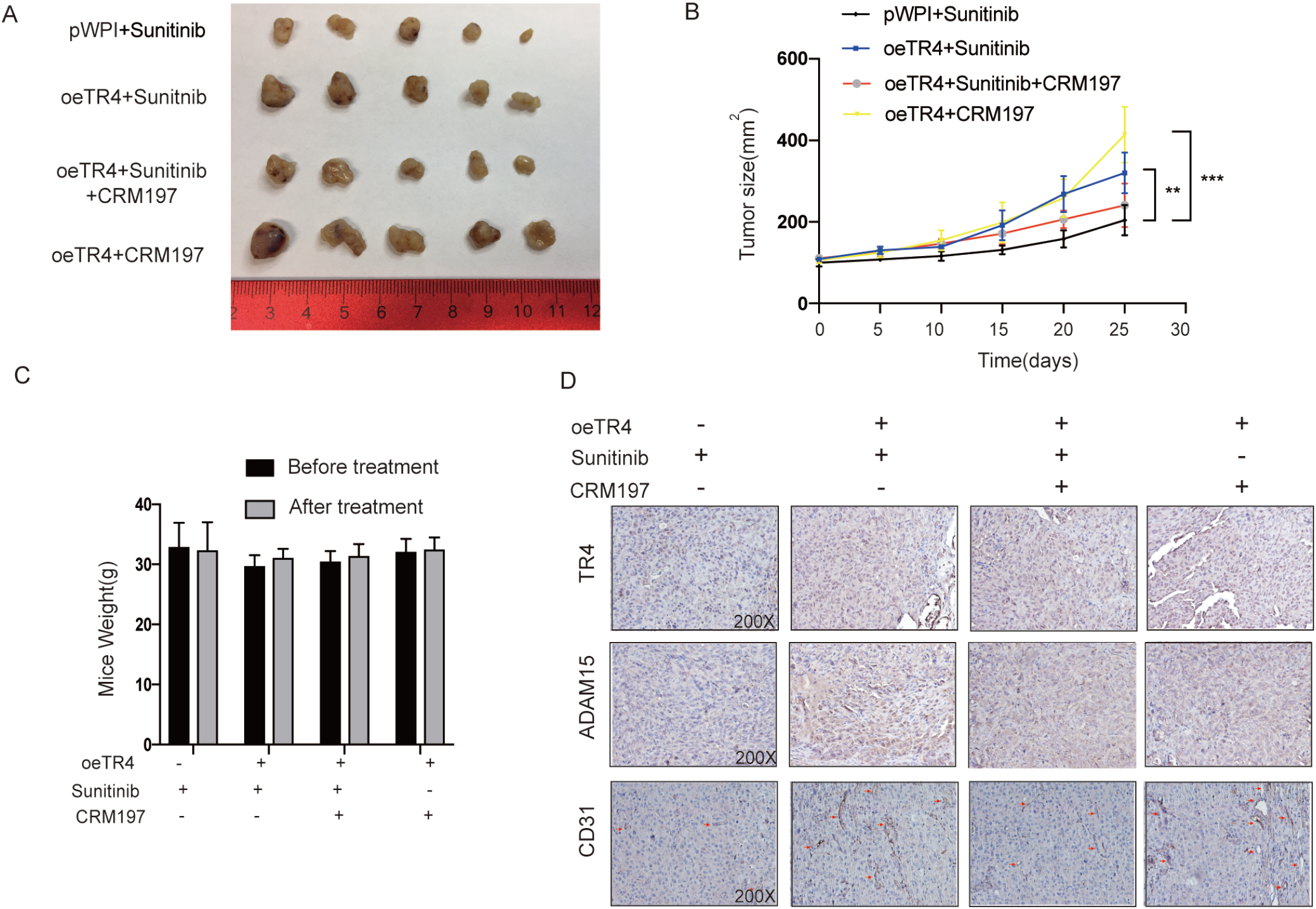
Targeting this newly identified signaling with CRM197,a HB-EGF inhibitor can enhance the sunitinib sensitivity of RCC in vivo. **A**. Representative isolated Tumors images from each treatment group at day25 after treatment are shown. **B**. Tumor size of the xenografts was observed in mice from each group. **C**. Mice weights were shown from each group after drug treatment. D. Positive expression of TR4, ADAM15 and CD31 proteins determined by immunohistochemistry (200 ×). E. A proposed model illustrating the role of the TR4/ADAM15/EGFR axis in regulating sunitinib sensitivity in this study. Data are presented as Mean ± SD.*P< 0.05, **P<0.01, ***P< 0.001, ****<0.0001. D. IHC staining of xenografts showing TR4, ADAM15 and CD31 stainings.

### TR4/ADAM15 axis can alter the angiogenesis in HUVECs via increasing the HB-EGF/EGFR signaling

Our finding that exosomes from RCC tumor cells with a higher TR4 expression can increase the EGFR phosphorylation while also increasing ADAM15 expression prompted us to test the hypothesis that ADAM15 can activate HB-EGF localized on the endothelial cell membrane through direct proteolytic cleavage followed by EGFR activation[36]. Indeed, early studies indicated that metalloproteases of the ADAM family were implicated as shedding enzymes of pro-HB-EGF for ErbB receptor activation[37].

To directly test that, we constructed an alkaline phosphatase AP-tag HB-EGF fusion gene as a probe to measure the ADAM15 activity in HUVEC tube formation assay (**Fig. 5A**). The cleavage of this fusion protein will result in an increase of alkaline phosphatase in the media, which can be measured as an approximation of the ADAM15 activity. The HUVECs transfected with the fusion gene construct were co-cultured with RCC cells followed by the alkaline phosphatase activity measurement in the CM(**Fig. 5B**). The results revealed that increasing RCC TR4 could increase AP activity in CM, and suppressing TR4 could decrease AP activity in CM. Importantly, oeTR4-increased AP activity could be partially blocked/reversed by suppressing ADAM15 **(Fig. 5C)**. As expected, the oeTR4-increased tube formation in HUVECs could be partially reversed *via* adding the HB-EGF neutralizing antibody in the co-culture system (**Fig. 5D**). Consistent with this, adding HB-EGF neutralizing antibody could also result in suppressing TR4-increased EGFR phosphorylation(**Fig. 5E**). Furthermore, CRM197, a specific HB-EGF inhibitor[38, 39], could partially reverse the oeTR4-increased tube formation in HUVEC cell and EGFR phosphorylation(**Fig. 5F-G**). Consistent finding was observed in OSRC-2 cells (**Fig. S2D**). Using CRM197 could reverse the oeTR4-increased tube formation in HUVEC cell.

The results confirmed the above findings, showing that the oeTR4-increased the quantity and length of vessels that sprouted from the aortic ring and could be partially reversed via adding the HB-EGF neutralizing antibody and CRM197**(Fig. 5H)**.

Together, results from **Fig. 5A-H** suggest TR4/ADAM15 axis may alter the angiogenesis/tube formation in HUVEC cells *via* modulating the HB-EGF/EGFR signaling.

### Targeting this newly identified signaling with CRM197, a HB-EGF inhibitor can enhance the sunitinib sensitivity of RCC in vivo

To test whether the mechanisms that we uncovered *in vitro* similarly play a significant role *in vivo*, we examined whether CRM197treatment can suppress RCC progression in a xenograft animal model. OSRC-2 cells with control or pwpi-oeTR4 were implanted subcutaneously into the right flank of immunodeficient nude mice. The RCC cells were divided into four groups, pWPI plus Sunitinib, oeTR4 with Sunitinib, oeTR4 plus Sunitinib and CRM197, oeTR4 plus CRM197. After 25 day’drug treatment, we sacrificed the mice and examined the tumor weights. Importantly, CRM197 can partly block/reverse the oeTR4-increased OSRC-2 cells growth under Sunitinib treatment (**Fig. 5A-B**), while on its own failed to suppress tumor growth. Mice weight were comparable among groups and CRM197 schedule did not lead to significant weight loss (**Fig.5C**). IHC were conduct to analysis the expression of ADAM15and CD31(vascular endothelial cell marker) in tumor. CRM197 can partly block/reverse the oeTR4-increased adam15 and microvessel density expression(**Fig. 5D**).

The above results indicated that the TR4-induced Renal cancer cells-derived exosomally initiated signaling with CRM197,a HB-EGF inhibitor can enhance the sunitinib sensitivity of RCC in vivo.

## Discussion

A large scale genomic analysis of the RCC clinical database from TCGA indicated that a higher TR4 expression results in a lower survival in RCC patients, but the detailed mechanisms underlying this finding are not clear. Recent studies indicated that TR4 could promote RCC VM formation to enhance RCC resistance to Sunitinib. Shi et al. also found that TR4 could promote RCC cells more resistant to Sunitinib[13]. Whether TR4 can also alter Sunitinib resistance *via* other mechanisms remains unclear.

Sunitinib as a multi-target inhibitor of tyrosine kinases is believed to be effective primarily through suppressing tumor angiogenesis as well as directly suppressing tumor proliferation. Resistance to Sunitinib in patients therefore needs to be investigated in these two critical aspects of drug efficacy. Indeed, its impact on suppressing tumor cell proliferation *in vitro* may not be approximating its effect *in vivo* as the concentration to suppress tumor cell survival far exceeds the available plasma concentration of Sunitinib(IC_50_ values required to inhibit RCC cell viability *in vitro* were in the range of 5μMwhile clinically relevant plasma concentrations of Sunitinib are only 0.1 to 0.2 μM[8]). Accordingly, our study attempted to examine the role of TR4 in Sunitinib efficacy through a more physiological setting of endothelial cells co-cultured with RCC cells. In addition, we also utilized a primary endothelial cells system with mouse aorta endothelial cell sprouting assay to test the potential role of TR4 in RCC cells and their impact on angiogenesis and sensitivity to Sunitinib in the form of endothelial cell tube formation. These studies were also verified using an*in vivo* mouse model with xenografts of RCC cells by targeting this newly identified signaling with the clinically available drug,CRM197, a specific HB-EGF inhibitor, which has been shown to significantly attenuate several tumor cells growth, including lung cancer[38], ovarian cancer[39], breast cancer[40]and glioblastoma cells[41]. We found that CRM197 can partly block/reverse the oeTR4-increased RCC tumor growth under Sunitinib treatment.

The solid tumors are regarded as “organs” composed of cancer cells and the tumor tumor microenvironment, including the extracellular matrix(ECM), mesenchymal stem cells(MSCs), endothelial cells and immune cells[42]. The connections between cancer cells and endothelial cells through the networks of cytokines[14], growth factors and exosomes[43] seems to be critical. Although studies concerning exosomes/microparticles, which can mediate intercellular exchange of bioinformation have been increasing, it is unclear how cancer cells can remodel endothelial cells *via* exosomes. In our study, we found that TR4 in renal cell lines increase angiogenesis and endothelial cell resistance to Sunitinib *via* altering the exosomes-mediated EGFR signal pathway in HUVEC.

ADAM15 has been shown to play a role in angiogenesis *via* producing the proinflammatory mediators, such as ENA78/CXCL5 and ICAM-1[25]. In addiction, ADAM15 is involved in the progression to metastatic diseases in several solid malignant tumors[44]. We found that silencing ADAM15 in RCC cells reversed TR4-induced angiogenesis of endothelial cells, which provides direct evidence supporting a role of ADAM15 in cancer cells’ promotion of tumor angiogenesis. As an ADAMs family member, which contains a zinc-binding metallopretease domain, ADAM15 has been implicated in the shedding of inflammatory cytokines and growth factors from the cell surface[25]. Consistent with this, we found that ADAM15 in HUVECs can induce endothelial cell tube formation through shedding HB-EGF and activating the EGFR signaling pathway.

EGFR signaling pathway activation has been shown to be associated with the stimulation of tumor angiogenesis. interestingly, the result also showed that EGFR phosphorylation was increased by overexpression of TR4 in RCC cells. It was attenuated by sunitinib with TR4-shRNA RCC cells, while it was augmented by sunitinib with TR4-cDNA RCC cells. We speculate that the increasing of EGFR phosphorylation need basal ADAM15. However, there are ADAM15 and inhibitor of EGFR phosphorylation. Without TR4, this inhibitor may prohibit EGFR phosphorylation. HB-EGF is an activating ligand for the EGF receptor (EGFR/ErbB1) and ErbB4. HB-EGF can induct HUVEC migration and capillary tube formation via activation of VEGF independent signal pathway which EGFR/PI3K/ERK1/2 signaling pathway[45]. To our knowledge, the role for TR4/ADAM15/HB-EGF in RCC angiogenesis during sunitinib treatment has not been previously demonstrated. In our study, we found that EGFR phosphorylation was increased by overexpression of TR4 in RCC cells. In addition, it was attenuated by sunitinib with TR4-shRNA RCC cells, while it was augmented by sunitinib with TR4-cDNA RCC cells. We speculate that sunitinib can regulate EGFR phosphorylation through ADAM15 which cleaves and releases HBEGF to act on endothelial cells. Indeed a previous report examining the VEGFR inhibitor sorafenib in HUVEC showed that vascular endothelial cells release s-HBEGF upon sorafenib stimulation[46]. When ADAM15 was induced by overexpression of TR4 in RCC cells, sunitinib resulted in more s-HBEGF to activate EGFR phosphorylation on endothelial cells. On the other hand, TR4-shRNA in RCC cells resulted in a lower ADAM15 thus a lower s-HBEGF while sunitinib inhibition of growth factor signaling in endothelial cells dominated with a reduction of EGFR phosphorylation.

## Conclusion

In summary, this study for the first time showed that TR4 in RCC blunted the efficacy of sunitinib via exosomal ADAM15-induced activation of EGFR signaling pathway in endothelial cells. Knowing down TR4 or blocking the TR4-induced renal cancers-derived exosomelly initiated signaling with a small molecular, CRM197, increases sunitinib efficacy in vitro and xenograft tumor models. It is therefore potentially feasible to determine whether exosomes-ADAM15 level can act as a gauge of TR4 expression as well as a diagnostic and prognostic marker for the efficacy of Sunitinib therapy for RCC patients. These mechanistic studies will help to develop new therapies to enhance Sunitinib efficacy for RCC.

## Materials and Methods Cell culture

The human normal cell line 293T and RCC cell lines 786-O and OSRC-2 were originally purchased from American Type Culture Collection (ATCC, Manassas, VA). The RCC cell line SW839 was purchased from Cell Resource Center for Biomedical Research, Tohoku University. All ccRCC cells were cultured in DMEM supplemented with 10% FBS in the humidified 5% CO2 environment at 37 °C. The vascular endothelium HUVEC-C cell line, which we use previously[27] was acquired from ATCC and maintained in media containing 10 ng/ml VEGF165 (Pepro Tech, USA). Cells were authenticated by STR typing and tested to have no mycoplasma or bacteria contamination before experiments.

### Tube formation assay

Endothelial cells (HUVEC) were co-cultured with RCC cells with indicated treatments (with/without overexpression/knockdown of TR4) for 72 h, with/without 1nMSunitinib for 24h and then the endothelial cells were harvested and subjected to the tube formation assay to measure migration and invasion capabilities. For these experiments, the endothelial cells were maintained in mixed conditioned media (CM) from the co-culture system with fresh media at the ratio of 1:1.

HUVEC cells co-cultured with SW839 cells with altered TR4 expression were trypsinized, then re-suspended in mixed CM at ∼2x104/ml. Growth factor-reduced Matrigel (BD Biosciences, San Jose, CA) at 50 μl were plated in 96-well plates at a horizontal level that allows the Matrigel to distribute evenly, and incubated for 1 h at 37°C. Then 100 μl of resuspended HUVECs were loaded onto the top of the matrigel. Each condition was performed at least in triplicate. Following the incubation at 37°C for 6 h, each well was analyzed under a microscope. Blinded quantification was done using Image J software to quantify tube lengths. Tube lengths in each field were imaged and numbers of tubules in 3-5 random fields for each well were counted and averaged.

### Mouse aortic ring assay

Matrigel was plated on Culture Slides (BD). C57BL/6mice(8-12 weeks) were sacrificed, and their thoracoabdominal aortas were procured. The aortas were dissected free of any fibroadipose tissue and sectioned into 0.5-mm rings. Rings were placed on Matrigel, one ring per chamber. Each ring was then embedded in Matrigel. Complete media was added, and the rings were incubated for another 24 h. The next day, media was exchanged for mixed CM with/without 0.01μMSunitinib. The rings were incubated in treatment media for 7 days, with media and drug compound being refreshed every other day; after 7 days, they were imaged on an inverted phase contrast microscope. Blinded quantification was done using Image J software to quantify tube lengths that represented the outgrowth. These pixel counts served as raw data for analysis.

### Construct RCC cell lines with differential stable expression of TR4

We knocked down and overexpressed TR4 in RCC cell lines according to our previous reports[11, 12].

### Quantitative real-time PCR

RNA isolation and quantitative RT-PCR were conducted according to our previous reports[11, 12]. Primers of TR4 and GAPDH were follows: TR4_F: 5′-TCC CCA CGC ATC CAG ATA ATC-3′; TR4_R: 5′-GAT GTG AAAACA CTC AAT GGG C-3′. GAPDH_F : 5′-GGA GCG AGA TCC CTC CAA AAT-3′; GAPDH_R: 5′-GGC TGT TGT CAT ACT TCT CAT GG-3′.

### Western blot analysis

Western blot were conducted according to our previous reports[11, 12].The primary antibodies used in the study for western blot were listed below: TR4 (Abcam, #ab109301, San Diego, CA),GAPDH (Santa Cruz, #sc-166574, Paso Robles, CA), ADAM15 (ABclonal, #A7756, Woburn MA), HB-EGF antibody (Novus Biologicals, #AF-259-Sp, Centennial, CO), p-EGFR (Y1068) (ABclonal, AP0301, Wuhan, CN), HB-EGF antibody (Novus Biologicals, #AF-259-Sp, Woburn, MA), EGFR (ABclonal, A11351, Wuhan, CN), IgG (Santa Cruz, #sc-2027, Dallas, TX).

### Luciferase reporter gene assays

Luciiferase reorter gene assays were conducted according to our previous reports[11, 12].The 1 kb human ADAM15 promoter was cloned into pGL3 basic vectors (Promega, Madison, WI).

### Chromatin Immunoprecipitation Assay (ChIP)

Normal IgG and protein A-agarose were used sequentially to preclear the cell lysates. We then added anti-TR4 antibody (2.0 μg) to the cell lysates and incubated overnight at 4 °C. IgG was used in the reaction for negative control. Specific primer sets were designed to amplify a target sequence within the human ADAM15 promoter and agarose gel electrophoresis was used to identify the PCR products.

### Establishent of xenograft mice models

The nude mice (age 6–8weeks) were purchased from the Animal Production Area of the National Cancer Institute-Fredrick Cancer Research and Development Center in Frederick and all the animal experiments were approved by performed in accordance with the National Institutes of Health Guide for the Care and Use of Laboratory Animals. The RCC cells OSRC were transfected with vectors, and subcutaneously injected into the back flanks of the mice for 14days. After that, the mice were sacrificed, the tumor weight was measured and the tissues were collected.

### H&E and immunohistochemical (IHC) staining

After sacrificing the mice, IHC was performed on the xenografted tumors. After fixing in 4% neutral buffered paraformaldehyde for 18 h, the samples were embedded in paraffin and then sequentially cut into 4 μm slices. After deparaffnization, hydration, antigen retrieval, and blocking, these slices were incubated with corresponding primary antibodies. The primary antibodies of the rabbit TR4 TR4 (Abcam, #ab109301, San Deigo, CA), the rabbit CD31 (ABclonal, #A7756, Woburn, MA) and ADAM15 (ABclonal, #A7756, Woburn, MA) were used for staining. then incubated with biotinylated secondary antibodies (Vector Laboratories, Burlingame, CA, USA), and finally visualized by VECTASTAIN ABC peroxidase system and 3, 3’-diaminobenzidine (DAB) kit (Vector Laboratories, Burlingame, CA). The slices were reviewed separately by two experienced pathologists. ImageJ software was applied to calculate the average optical density of staining.

### Statistics

All experiments were performed in triplicate, and the results are presented as the mean value ± standard deviation. The data were statistically analyzed using ANOVA. Student’s t-test in SPSS statistical software, with P < 0.05 considered statistically significant. * indicates P < 0.05; ** indicates P < 0.01 and *** indicates P < 0.001.

## Abbreviations

RCC: Renal cell carcinoma
TR4: Testicular orphan receptor 4
ADAM15: A Disintegrin and Metalloproteinase 15
EGFR: epidermal growth factor receptor
HB-EGF: Heparin-binding EGF-like growth factor
HC: Immunohistochemistry
ChIP: Chromatin immunoprecipitation
WB: Western blot.

## Declarations

### Ethics approval and consent to participate

All study participants provided informed consent, and the study design was approved by the appropriate ethics review board.

## Acknowledgements

Not applicable.

## Authors’ contributions

ZWW, YS, XFQ GSY and CSC designed the experimental protocols. ZWW, HYX, CPH, FJC and BSY performed the studies. ZWW and HYX analyzed the data. ZWW, YS, XFQ, GSY and CSC wrote the manuscript with contributions from all of the other authors.

## Availability of data and materials

All data generated or analyzed during this study are included in this published article and its supplementary information files.

## Consent for publication

Not applicable.

---

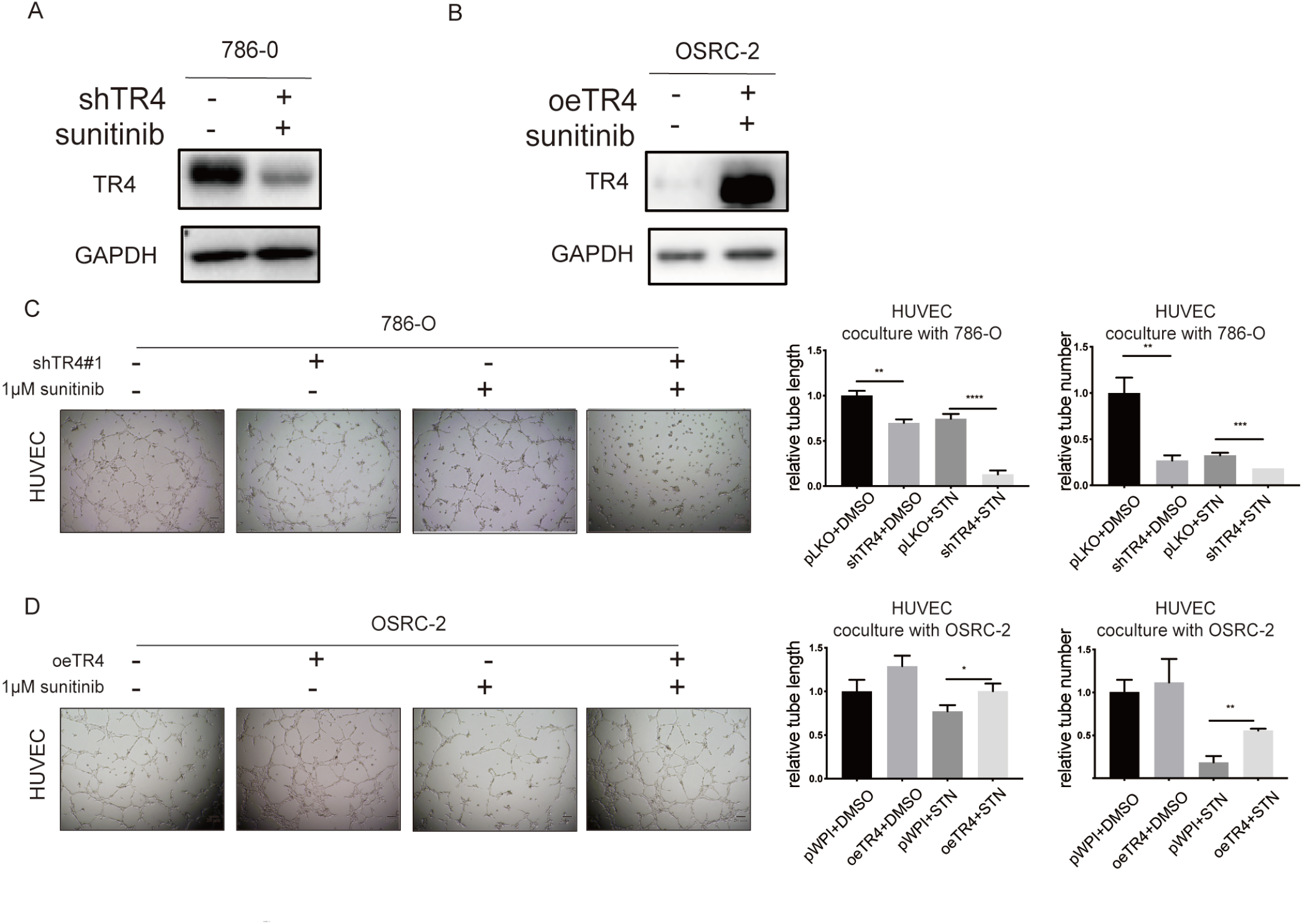

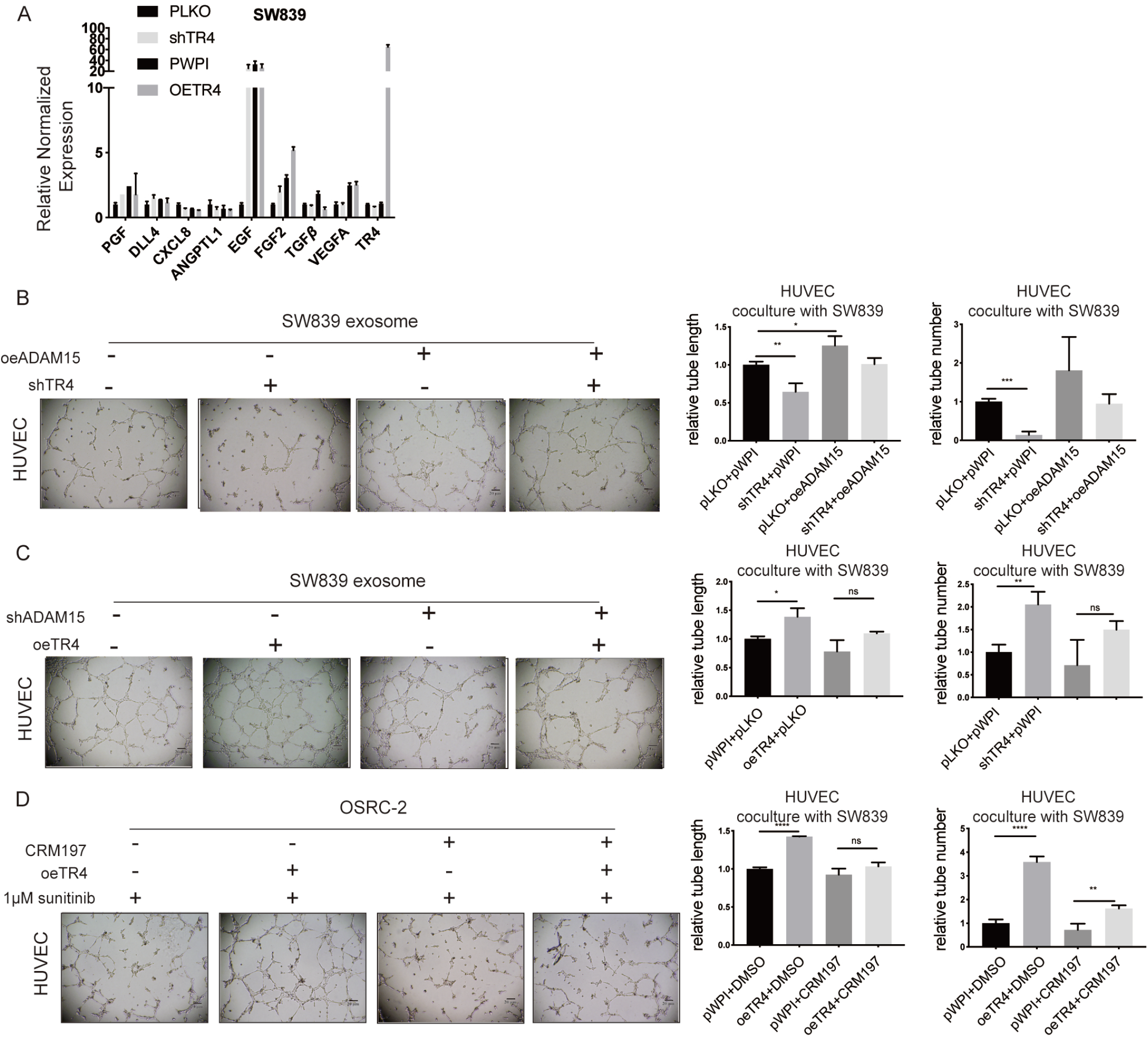

